# Neural Signatures of α2 Adrenergic Agonist-Induced Unconsciousness and Awakening by Antagonist

**DOI:** 10.1101/2020.04.21.053330

**Authors:** Jesus J. Ballesteros, Jessica Briscoe, Yumiko Ishizawa

## Abstract

How the brain dynamics transition during anesthetic-induced altered states of consciousness is not completely understood. The *α*2 adrenergic agonist is a unique anesthetic that generates unconsciousness selectively through *α*2 adrenergic receptors and related circuits. We studied intracortical neuronal dynamics during transitions of loss of consciousness (LOC) with the *α*2 adrenergic agonist dexmedetomidine and return of consciousness (ROC) in a functionally interconnecting somatosensory and ventral premotor network in non-human primates. LOC, ROC and full task performance recovery were all associated with distinct neural changes. The early recovery demonstrated characteristic intermediate dynamics distinguished by sustained high spindle activities. Awakening by the *α*2 adrenergic antagonist completely eliminated this intermediate state and instantaneously restored awake dynamics and the top task performance while the anesthetic was still being infused. The results suggest that instantaneous functional recovery is possible following anesthetic-induced unconsciousness and the intermediate recovery state is not a necessary path for the brain recovery.

## Introduction

General anesthetics are unique drugs that can induce reversible unconsciousness and have been widely used for over 170 years. However, the neurophysiological mechanisms of anesthetic-induced altered states of consciousness are not completely understood. Recent electroencephalography (EEG) and neuroimaging studies significantly advanced our understanding of the brain activity during general anesthesia. Anesthetic-induced unconsciousness is now thought to be associated with profound oscillations between brain structures (Lewis *et al.*, 2012; Purdon *et al.*, 2013; Akeju *et al.*, 2014b; Ishizawa *et al.*, 2016; Ballesteros *et al.*, 2020; Patel *et al.*, 2020). Moreover, we recently reported that anesthetic-induced state transitions are abrupt during propofol-induced loss of consciousness (LOC) and return of consciousness (ROC) in non-human primates (Ishizawa *et al.*, 2016; Patel *et al.*, 2020). Discrete metastable states have been reported during emergence from isoflurane in small animals (Hudson *et al.*, 2014). Abrupt state transitions might be a fundamental manner of how the brain functions during anesthetic-induced altered states of consciousness. However, oscillatory dynamics per se during transitions and unconsciousness appear to be unique to each anesthetic agent.

Dexmedetomidine is a highly selective α2 adrenergic agonist and is unique among currently available anesthetics, most of which are known to act at multiple receptors in the central nervous system. A recent neuroimaging study suggests that dexmedetomidine significantly reduces information transfer in the local and global brain networks in humans (Hashmi *et al.*, 2017), consistent with the effect of propofol (Monti *et al.*, 2013). Functional disconnection between brain regions is a possible mechanism suggested for dexmedetomidine-induced unconsciousness (Akeju *et al.*, 2014a; Song *et al.*, 2017). EEG changes under dexmedetomidine anesthesia are distinctive, with increased slow-delta oscillations across the scalp and increased frontal spindle oscillations, and closely approximate the dynamics during human non-rapid eye movement (NREM) sleep (Akeju *et al.*, 2014b; Akeju *et al.*, 2016). However, direct recordings from neocortex, especially from functionally interconnected regions, with α2 adrenergic agonist are rare, despite its increasing role in clinical anesthesia. Moreover, dexmedetomidine is unique because a specific α2 adrenergic antagonist is used in veterinary medicine to reverse its sedative and anesthetic effects (Scheinin *et al.*, 1998; Kamibayashi and Maze, 2000). Neuronal dynamics of awakening by the α2 adrenergic antagonist need to be studied.

Here we investigated how neural dynamics change during dexmedetomidine-induced LOC and ROC by directly recording from a functionally and anatomically interconnecting somatosensory (S1) and ventral premotor area (PMv) network, a well-studied corticocortical circuit, in non-human primates (**Fig. 1A**) (Kurata, 1991; Tanne-Gariepy *et al.*, 2002; de Lafuente and Romo, 2006; Garbarini *et al.*, 2019). The PMv is known to link sensation and decision-making as well as to integrate multisensory modalities (Rizzolatti *et al.*, 2002; de Lafuente and Romo, 2005, 2006; Pardo-Vazquez *et al.*, 2008; Lemus *et al.*, 2009; Acuna *et al.*, 2010; Romo and de Lafuente, 2013). We recorded local field potentials (LFPs) and single unit activity using surgically implanted microelectrode arrays in two adult macaque monkeys during dexmedetomidine-induced LOC and throughout recovery. We defined LOC and two recovery endpoints, ROC and return of preanesthetic performance level (ROPAP), using a probability of task engagement and task performance (Patel *et al.*, 2020) (**Fig. 1B, C**). Task engagement indicates a probability of any response initiation, including correct responses and failed attempts, and task performance represents a probability of correct responses only (Wong *et al.*, 2011; Wong *et al.*, 2014) (**Fig. 1B**). Dexmedetomidine was infused through a surgically implanted vascular port. We further investigated how the α2 adrenergic antagonist effectively reverse dexmedetomidine-induced unconsciousness.

**Figure 1.**
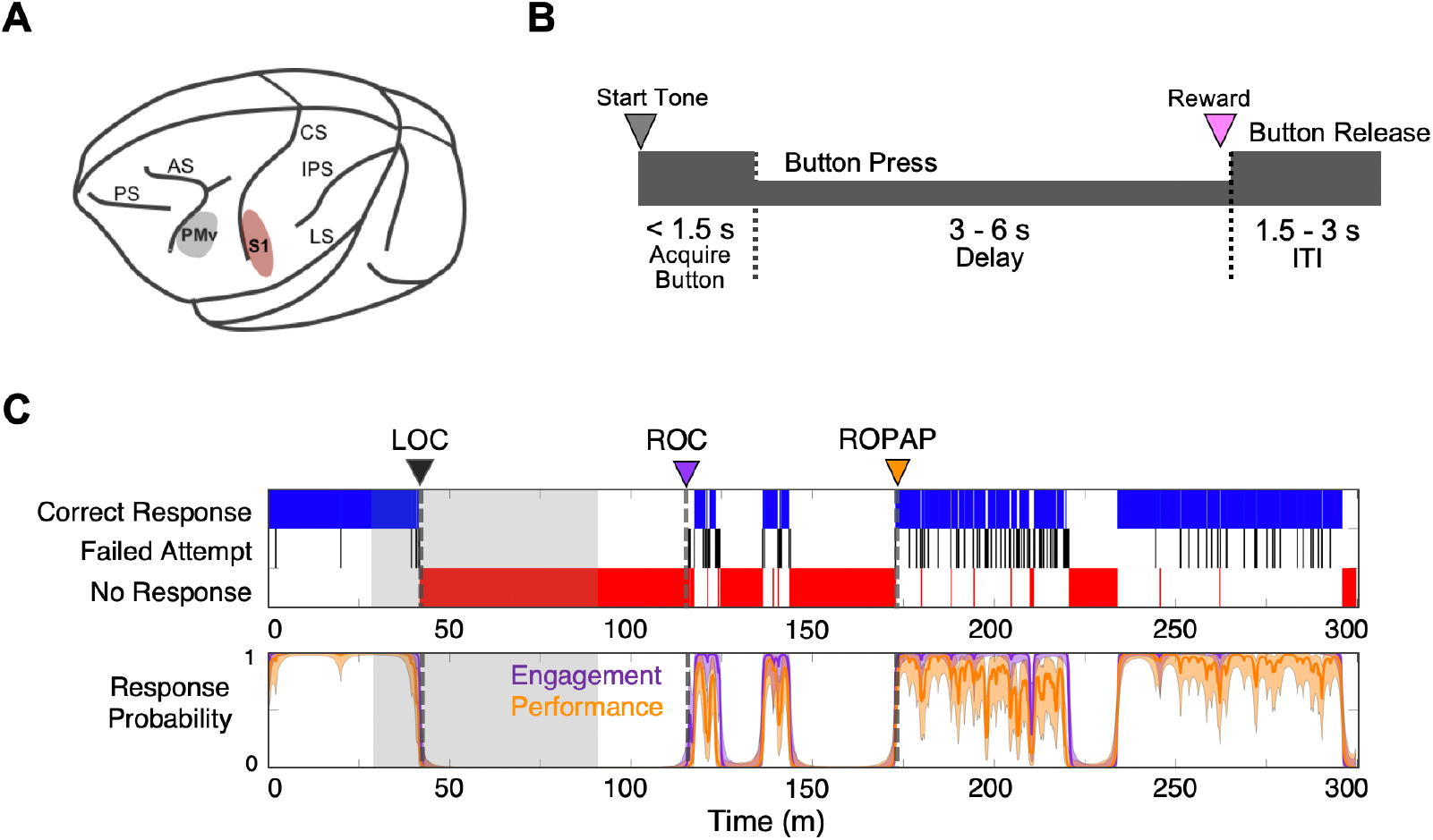
Experimental paradigm and behavioral responses. **A. Location of the recording sites.** Neural recording is performed in the primary somatosensory cortex (S1, red) and ventral premotor cortex (PMv, gray). CS, central sulcus; IPS, intraparietal sulcus; LS, lateral sulcus; AS, acuate sulcus, and PS, principal sulcus. **B. Behavioral task.** Sequence of events during behavioral trials. After the start tone (pure tone 1000 Hz 100 msec), the monkey initiates a trial by placing the hand (ipsilateral to the recording site) on the button in front of the animal. The animal is required to keep its hand on the button until the end of the trial in order to receive a liquid reward (correct response). The animal then has to release the button during the inter-trial interval (ITI). **C. Typical behavioral response during dexmedetomidine induction and emergence**. Following the start of dexmedetomidine infusion, failed attempts (black) increased briefly before the animal completely lost the response. Top: The animal’s trial-by-trial button response. Correct responses (blue), failed attempts (black), and no response (red). Bottom: Probability of the task engagement (correct and failed responses, purple) and task performance (correct response only, orange). LOC was defined as the time at which the probability of task engagement was decreased to less than 0.3, and ROC was defined as the first time, since being unconscious, at which the probability of task engagement was greater than 0.3. ROPAP was defined as the time at which the probability of task performance was returning to greater than 0.9 since being unconscious and remained so for at least 3 minutes. LOC is shown with a black arrow and dotted lines, ROC with a purple arrow and dotted lines, and ROPAP with an orange arrow and dotted lines (**C**). Dexmedetomidine was infused at 18 μg/kg/h for the first 10 min and then 4 μg/kg/h for 50 min (shaded area in **C**).

## Results

We successfully determined LOC, ROC and ROPAP in all the recording sessions in both animals. Interestingly, the task response behaviors appeared fluctuating, and a performing period and a non-performing period were often rapidly alternating over the course of recovery, especially during early recovery following ROC, as shown in **Figure 1A** and **Figure 2A**.

**Figure 2.**
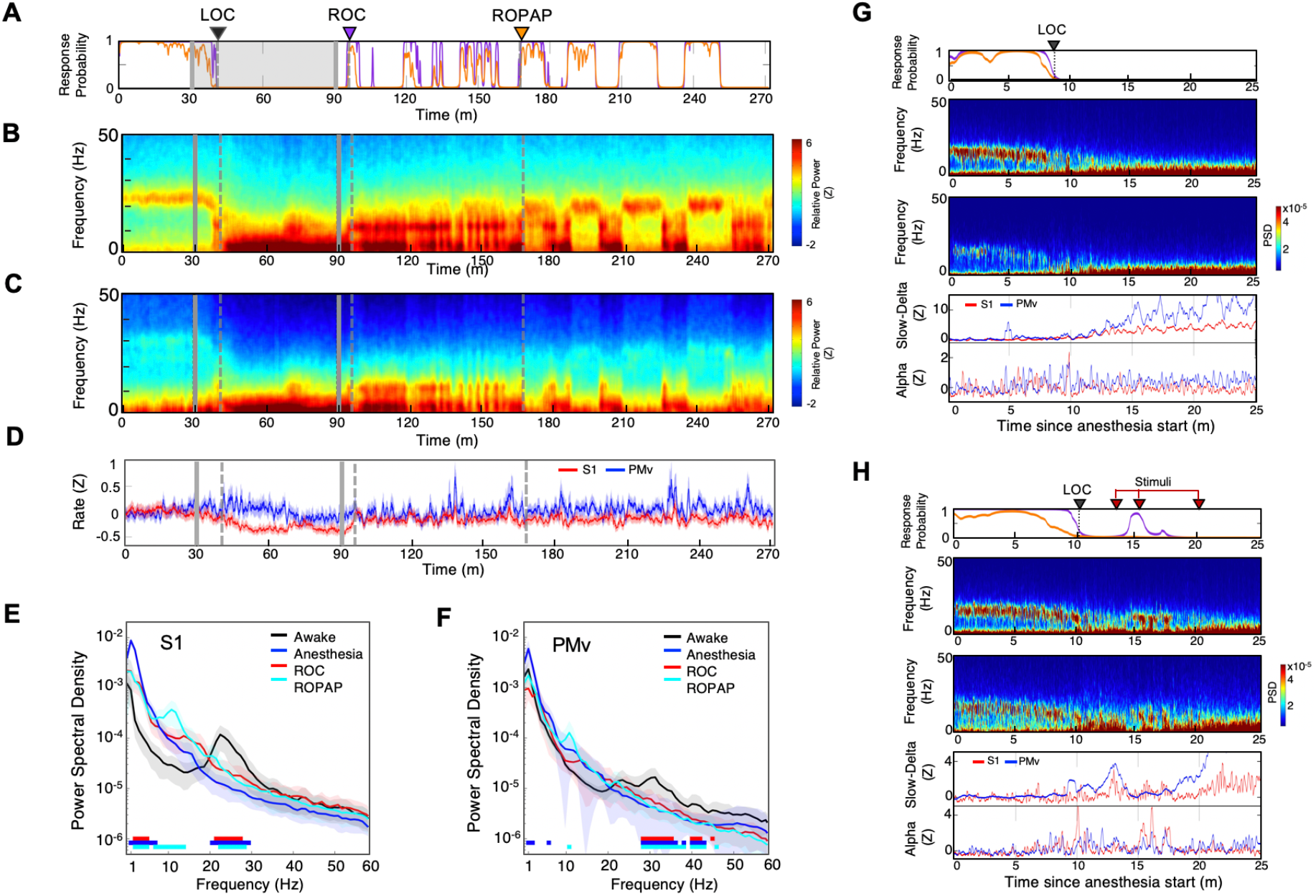
Distinctive neural changes are associated with α2 adrenergic agonist-induced LOC, ROC and ROPAP. **A.** Behavioral response. Probability of the task engagement (purple) and task performance (orange). **B.** Local field potentials (LFP) time-domain spectrograms in S1. **C.** LFP time-domain spectrograms in PMv. **D.** Average baseline firing rates in S1 (red) and PMv (blue). Firing rates were normalized to pre-anesthetic values using Z-scores. No significant differences between pre-anesthetic average and any following time point were found (ANOVA and *post hoc* Bonferroni multiple comparison tests). **E, F.** Averaged frequency domain power spectra in S1 (**E**) and PMv (**F**). Traces are the averaged Welch’s power across channels with 95% confidence intervals shaded, during wakefulness (the last minute before anesthesia infusion, black), anesthesia (the last minute of dexmedetomidine infusion, blue), ROC (first minute after ROC, red) and ROPAP (first minute after ROPAP, cyan). Bottom lines represent those frequencies with significantly different values of average power density between awake and any other given condition (ANOVA and *post hoc* Bonferroni multiple comparison test, p-value < 0.05). **G.** Behavioral response, spectrogram, the slow-delta (0.5-4 Hz) and alpha power (8-12 Hz) change during LOC. Power was normalized to pre-anesthetic values using Z-scores. **H.** Behavioral response, spectrogram, the slow-delta (0.5-4 Hz) and alpha power (8-12 Hz) change during LOC with arousability testing. A series of non-aversive stimuli (ear-pulling, a loud white noise at 100 dB SPL for 5 sec, and hand claps 3 times at 10 cm from face, shown with red arrows) were applied at 3, 5, and 10 min after initially detected LOC. LOC is shown with a black arrow and dotted lines, ROC with a purple arrow and dotted lines, and ROPAP with an orange arrow and dotted lines (**A-C**). Dexmedetomidine was infused at 18 μg/kg/h for the first 10 min and then 4 μg/kg/h for 50 min (gray lines in **A-C**).

### Distinctive neural changes at α2 adrenergic agonist-Induced LOC, ROC and ROPAP

We first compared LFP spectrograms from the primary somatosensory cortex (S1) and ventral premotor area (PMv) during the transition from wakefulness to LOC and then through recovery. During wakefulness beta oscillations were present in both cortical regions (18-25Hz in S1, 26-34Hz in PMv, **Fig. 2B, C**) (Brovelli *et al.*, 2004; Haegens *et al.*, 2011). The LOC was identified at a brief increase of the alpha power following disruption of the beta oscillations. Then the slow-delta oscillations appeared and remained dominant throughout anesthesia until ROC. ROC was associated with an abrupt diminishing of the slow-delta oscillations and an appearance of alpha oscillations (**Fig.2B, C**). ROPAP was observed at return of the beta oscillations. The peak frequencies of the beta oscillations, however, appeared to remain significantly lower than that during wakefulness (**Fig.2E, F**). During an early recovery period following ROC, the slow-delta oscillations returned repeatedly when the animal was not engaged in the task and appeared to be coupled with the alpha oscillations. Both slow-delta and alpha oscillations disappeared when the beta activity returned, suggesting that two or more states were alternating until full functional recovery. We also found that dexmedetomidine did not significantly change the average firing rate in the S1 units nor in the PMv units (**Fig. 2D**).

We then tested arousability during the period following LOC using a short series of non-aversive stimuli in separate sessions. Contrary to a regular session without stimuli (**Fig. 2G**), we observed a brief return of task attempts upon the stimuli at 3 minutes and 5 minutes after initially detected LOC (**Fig. 2H**). The slow-delta oscillations were disrupted, and a brief reappearance of alpha oscillations were observed in the spectrogram when the animal’s task attempts returned upon the stimuli, suggesting that the slow-delta oscillations per se do not assure non-arousable state. Arousability was not observed by the same stimuli at 10 minutes after the initial LOC and there was no change in the oscillatory dynamics (**Fig. 2H**).

We next investigated how dexmedetomidine affects communication across S1 and PMv by examining both local and regional coherence changes. We found that beta oscillations were strongly coherent between S1 and PMv during awake task performance (**Fig. 3A-F**). However, following the start of dexmedetomidine infusion, while the animal was still performing the task, coherent beta oscillations appeared to be disrupted locally and inter-regionally. Subsequent dynamics characterized by a brief appearance of alpha oscillations at LOC and evolving slow-delta oscillations seemed to be all coherent locally and inter-regionally. During recovery, the alpha and slow-delta oscillations and the beta oscillations were also coherent inter-regionally, suggesting the network is coherent throughout anesthesia and recovery. The beta oscillations at ROPAP appeared to be equally coherent inter-regionally to those during wakefulness, but the peak beta frequencies at ROPAP were still significantly lower than the awake level (**Fig. 3G**).

**Figure 3.**
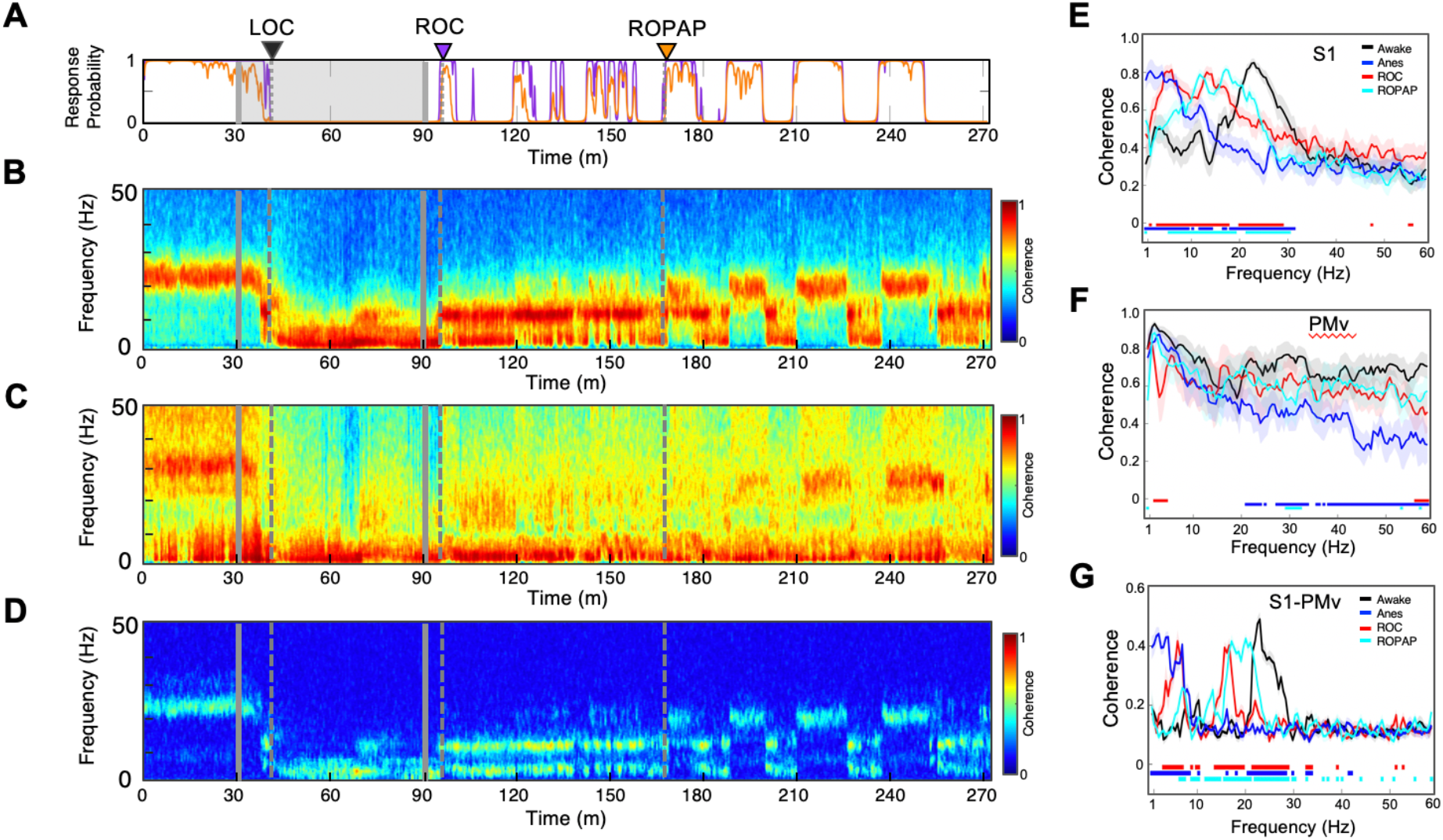
Oscillatory dynamics are inter-regionally coherent through α2 adrenergic agonist-induced anesthesia and recovery. **A.** Behavioral response. Probability of the task engagement (purple) and task performance (orange). **B.** Local field potentials (LFP) time-domain coherogram in S1. **C.** LFP time-domain coherogram in PMv. **D.** LFP time-dime coherogram between S1 and PMv. **E-G.** Averaged frequency-domain coherence within S1 (**E**), within PMv (**F**), and between S1 and PMv (**G**) at awake (for one minute before anesthesia start, black), anesthesia (for one minute before the end of anesthesia infusion, blue), ROC (for one minute after ROC, red) and ROPAP (for one minute after ROPAP, cyan). Coherence was averaged for a 1-minute period across all pair-wise channel for each epoch. Average lines are shown with shaded 95% confidence intervals. Bottom lines represent those frequencies with significantly different values of average coherence between awake and any other given condition (ANOVA and *post hoc* Bonferroni multiple comparison test, p-value < 0.05). LOC is shown with a black arrow and dotted lines, ROC with a purple arrow and dotted lines, and ROPAP with an orange arrow and dotted lines (**A-D**). Dexmedetomidine was infused at 18 μg/kg/h for the first 10 min and then 4 μg/kg/h for 50 min (gray lines in **A-D**).

There was no comparable neurophysiological change at the loss of response or at the return of response in alert behaving animals without anesthetic (Ishizawa *et al.*, 2016), suggesting that behavioral changes due to satiety or motivation are unlikely to be associated with the neurophysiological changes observed during dexmedetomidine-induced LOC or ROC.

### Spindle activity was highest during an early recovery period

We also investigated spindle oscillations during dexmedetomidine anesthesia and recovery. Spindle density was analyzed in the alpha and low beta frequencies between 9 and 17 Hz over the course of behavioral changes with dexmedetomidine (Kam *et al.*, 2019) (**Fig. 4A, B**). We found that spindle activity emerged prior to LOC and the spindle density initially peaked at LOC. Spindle activity was present through dexmedetomidine-induced unconsciousness, and then further increased upon the end of anesthetic infusion and through ROC (**Fig. 4B**). Spindle activity remained higher when the animal was performing the task prior to ROPAP, as compared to the performing period after ROPAP where the spindles were nearly completely diminished. Moreover, the spindle activity appeared to increase at the behavioral transitions between task responding and non-responding periods. The power of alpha frequencies seemed to be correlating with the spindle activity during anesthesia and recovery (**Fig. 4C**). Spindle characteristics, including density, duration and peak frequency, were similar between under anesthesia and during a non-performing period after ROPAP, but spindle density was statistically significantly higher during the non-performing period after ROPAP in S1 and the duration was longer in the non-performing period in S1 and PMv (**Fig. 4D, E, F**).

**Figure 4.**
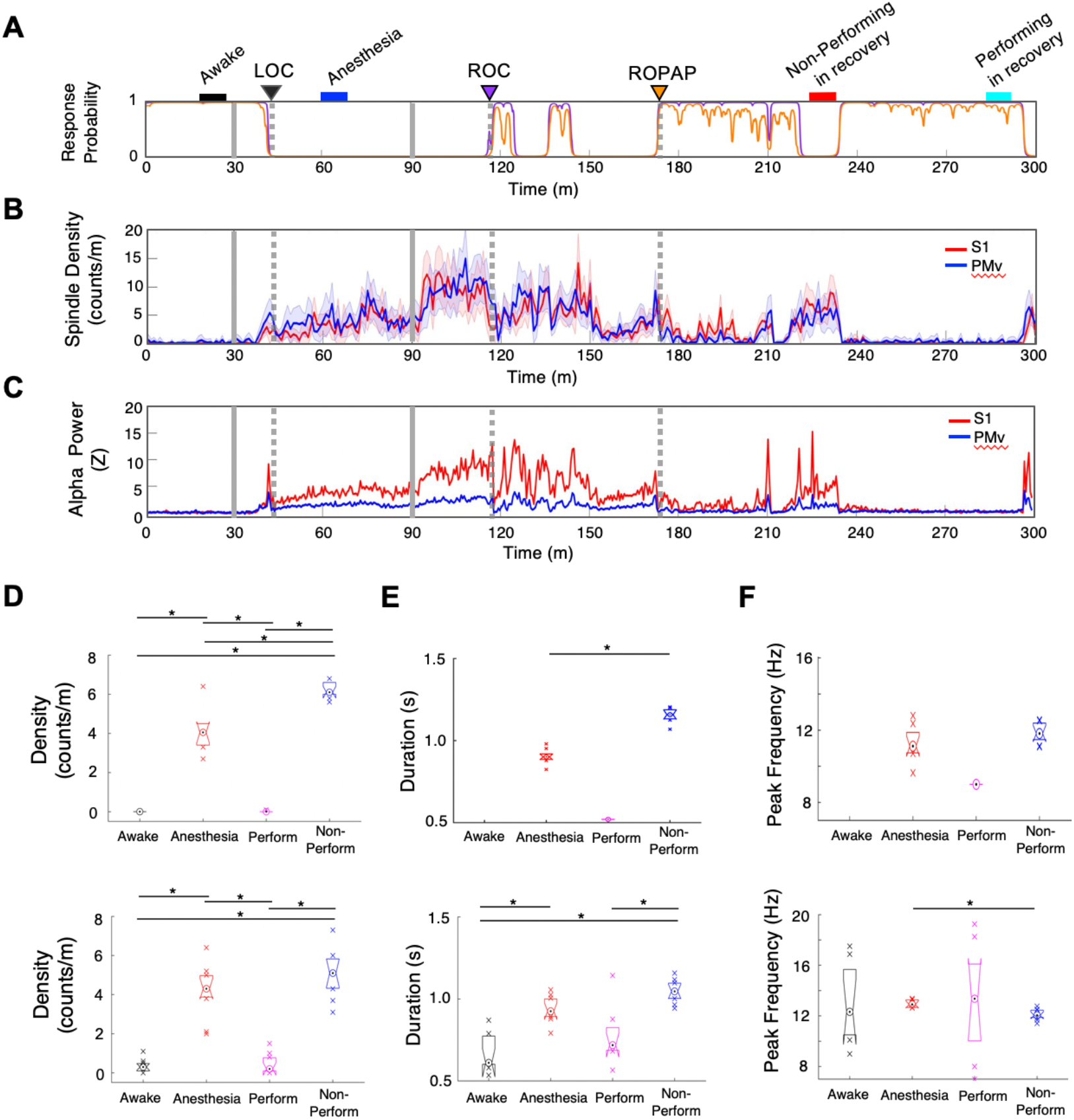
Spindle activity increases during early recovery following α2 adrenergic agonist-induced anesthesia. **A.** Behavioral response. Probability of the task engagement (purple) and task performance (orange). **B.** Spindle density (counts/min) in 9–17 Hz in S1 (red trace) and PMv (blue trace). **C.** Normalized power of alpha frequency (8-12 Hz) in S1 (red trace) and PMv (blue trace). **D-F.** Spindle characteristics in S1 (top plots) and PMv (bottom plots) of median density (**D**), median duration (**E**), and median peak frequency (**F**). Comparisons were made between awake (for 10 minutes before anesthesia start, black bar in **A**), anesthesia (for 10 minutes around mid-infusion, red bar in **A**), performing after ROPAP (for the last 10 minutes of the performing period, blue bar in **A**) and non-performing after ROPAP (for 10 minutes of the non-performing period, pink bar in **A**). Box plots represent the median with 25^th^ and 75^th^ percentiles and single points are all values beyond. Asterisks indicate statistically significant difference (two-sided unpaired t-test, \**p* < 0.01). LOC is shown with a black arrow and dotted lines, ROC with a purple arrow and dotted lines, and ROPAP with an orange arrow and dotted lines (**A-C**). Dexmedetomidine was infused at 18 μg/kg/h for the first 10 min and then 4 μg/kg/h for 50 min (gray solid lines in **A-C**).

### *α*2 adrenergic antagonist immediately restored awake dynamics and top task performance without intermediate stages

*α*2 adrenergic antagonist atipamezole induced an instant return of the top task performance while dexmedetomidine was still being infused (**Fig. 5A**). Concurrently, oscillatory dynamics demonstrated discontinuous return of the robust beta oscillations at the frequencies that were shown during wakefulness (**Fig. 5B, C, F, G**). These beta oscillations were inter-regionally coherent (**Fig. 5D, H**). In fact, the line coherogram indicated that the beta oscillations were significantly more coherent inter-regionally after *α*2 antagonist administration than that during wakefulness (**Fig. 5G**). The spindle activity was abruptly diminished upon the antagonist administration without its increase prior to ROC that was observed during recovery without antagonist.

**Figure 5.**
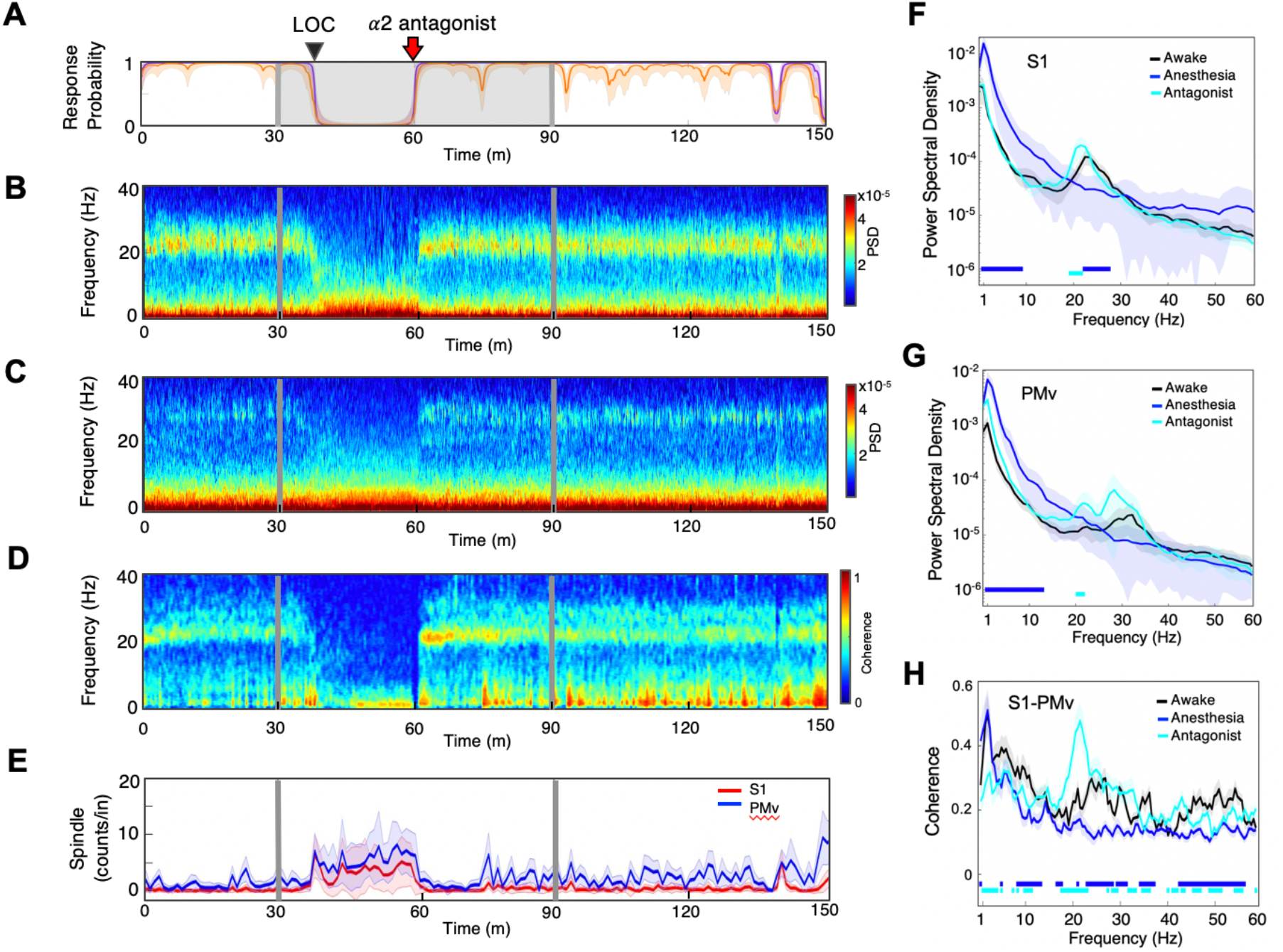
α2 adrenergic antagonist immediately restores top task performance and awake dynamics. **A.** Behavioral response. Probability of the task engagement (purple) and task performance (orange). **B.** Local field potentials (LFP) time-domain spectrogram in S1. **C.** LFP time-domain spectrogram in PMv. **D.** LFP time-domain coherogram between S1 and PMv. **E.** Spindle density (counts/min) in 9–17 Hz in S1 (red trace) and PMv (blue trace). **F-G.** Averaged frequency-domain power spectra in S1 (**F**) and PMv (**G**). Traces are the averaged Welch’s power across channels with shaded 95% confidence intervals, during wakefulness (for the last minute before anesthesia start, black), anesthesia (for the last minute of dexmedetomidine infusion, blue) and immediately following α2 adrenergic antagonist injection (Antagonist, cyan). Bottom lines represent those frequencies with significantly different values of average power density between awake and any other given condition (ANOVA and *post hoc* Bonferroni multiple comparison test, p-value<0.05). **H.** Averaged frequency-domain coherence between S1 and PMv at awake (for the last minute before anesthesia start, black), anesthesia (for the last minute of dexmedetomidine infusion, blue) and immediately following α2 adrenergic antagonist injection (Antagonist, cyan). Average lines are shown with shaded 95% confidence intervals. Bottom lines represent those frequencies with significantly different values of average coherence between awake and any other given condition (ANOVA and *post hoc* Bonferroni multiple comparison test, p-value<0.05). LOC is shown with a black arrow and dotted lines (**A-D**). Dexmedetomidine was infused at 18 μg/kg/h for the first 10 min and then 4 μg/kg/h for 50 min (gray lines and shaded area in **A-D**). α2 adrenergic antagonist atipamezole 100 μg/kg was intravenously injected after 30 minutes of dexmedetomidine infusion (at the time of 60 minutes).

Focusing on the state transitions, we investigated the dynamics in a three-dimensional (3D) state space to further characterize the transitional dynamics during recovery without *α*2 antagonist (**Fig. 6A, C**) versus with *α*2 antagonist (**Fig. 6B, D**) (Gervasoni *et al.*, 2004; Hudson *et al.*, 2014). During recovery without antagonist, we found two distinct clusters with a high-density core and a number of clouds connecting these two clusters in both S1 and PMv, possibly forming an intermediate cluster (**Fig. 6A_1, 2_, C_1, 2_**). The intermediate area corresponded to high speed values of spontaneous trajectories (**Fig. 6A_3_, C_3_**) and mixed task responses (**Fig. 6A_4_, C_4_**), indicating unstable transitions. Two distinct clusters appeared to be distinguished by the animal’s task performance level, high task performance versus minimum - zero task response. Awakening induced by *α*2 antagonist atipamezole demonstrated no intermediate state and two clusters were clearly separated in the state space in S1 nor in PMv (**Fig. 6B_1, 2_, D_1, 2_**). The state transition from unresponsiveness to responsiveness was discontinuous and abrupt (**Fig. 6B_3_, D_3_**). Two clusters representing unconsciousness and wakefulness were exclusively associated with no response and the highest performance, respectively (**Fig. 6B_4_, D_4_**).

**Figure 6.**
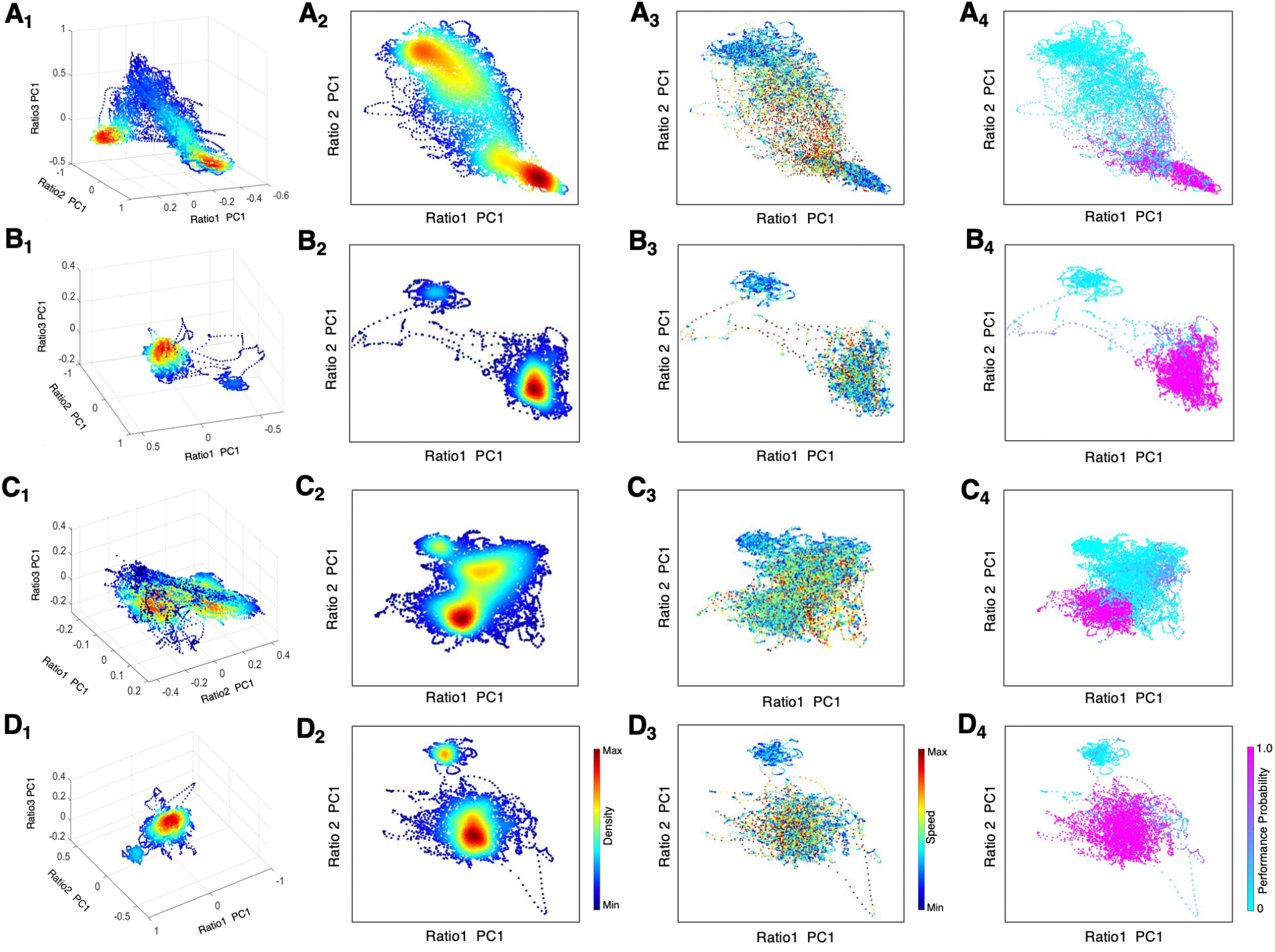
State space characterizes recovery through intermediate state without α2 antagonist and instant awakening with α2 antagonist. **a_1_-d_1_. 3D density plots** during anesthesia and recovery without antagonist in S1 (**A_1_**) and PMv (**C_1_**) and with antagonist in S1 (**B_1_**) and PMv (**D_1_**). The density heat-map was calculated with a kernel density estimator from the scatter data created using the first principal component from the three spectral ratios described in the methods. **A_2_-D_2_. Density plots in 2D** during anesthesia and recovery without antagonist in S1 (**A_2_**) and PMv (**C_2_**) and with antagonist in S1 (**B_2_**) and PMv (**D_2_**). **A_3_-D_3_. Speed plots** during anesthesia and recovery without antagonist in S1 (**A_3_**) and PMv (**C_3_**) and with antagonist in S1 (**B_3_**) and PMv (**D_3_**). The speed was calculated as the Euclidean distance between consecutive points and is represented as heat-map. **A_4_-D_4_. Task performance** during anesthesia and recovery without antagonist in S1 (**A_4_**) and PMv (**C_4_**) and with antagonist in S1 (**B_4_**) and PMv (**D_4_**). The plots are color-coded according to the task performance probability. The data were analyzed for the period between LOC and the end of recording session. Each dot corresponds to a 1-second window for all plots. Dexmedetomidine was infused at 18 μg/kg/h for the first 10 min and then 4 μg/kg/h for 50 min in all sessions. α2 adrenergic antagonist atipamezole 100 μg/kg was intravenously injected at 30 minutes of dexmedetomidine infusion.

## Discussion

Our results demonstrate that distinctive, not gradual, neural changes were associated with the behavioral endpoints during *α*2 adrenergic agonist dexmedetomidine-induced altered states of consciousness, consistent with propofol-induced LOC and ROC (Ishizawa *et al.*, 2016; Patel *et al.*, 2020), even though a pharmacokinetic model assures gradual change in the anesthetic concentrations (Patel *et al.*, 2020). Abrupt or non-linear state transitions are well known during natural sleep (Saper *et al.*, 2010; Stevner *et al.*, 2019) and in epilepsy (Bartolomei and Naccache, 2011). Together, these results suggest that abrupt state transitions are a fundamental manner of how the brain functions.

In the current work, a brief appearance of alpha oscillations with an increase of spindle activity were characteristic at LOC and ROC, and the slow-delta oscillations became dominant after LOC and through ROC. Full task performance recovery also appeared to be associated with return of robust beta oscillations. Additionally, these characteristic oscillations were coherent locally as well as inter-regionally during anesthesia and recovery, suggesting that dexmedetomidine does not interrupt consistency of the dynamics in this cortical network. Interestingly, we have shown discrepancy between local coherence and inter-regional coherence during propofol anesthesia (Ishizawa *et al.*, 2016). Especially, the high beta-gamma oscillations at propofol-induced LOC that were highly coherent locally were not coherent inter-regionally. Together, these results suggest that dexmedetomidine preserves the continuity across frequencies through the state changes while propofol interrupts cortico-cortical communication during the state transition. The results also suggest that the abrupt neural changes in the cortical dynamics induced by *α*2 adrenergic agonist are likely driven by remote areas, such as subcortical regions.

Recovery from dexmedetomidine shows a characteristic intermediate state with an increase in the alpha power and high spindle density, especially during early recovery following the end of anesthetic infusion through ROC. Oscillatory dynamics of this intermediate state resembles the state of NREM and REM sleep (Prerau *et al.*, 2017) and is finally replaced with coherent beta oscillations characteristic to wakefulness. Zhang and colleagues reported that sleep-like sedation with *α*2 adrenergic agonist is mediated through the preoptic hypothalamic area, while loss of righting reflex, considered a surrogate for unconsciousness, requires locus coeruleus (LC) using pharmacogenetic techniques in mice (Zhang *et al.*, 2015). Observed behavioral state changes under dexmedetomidine in the current study, including unresponsiveness and an intermediate state with fluctuating task responses, are consistent with their findings of neuroanatomical sites of the dexmedetomidine action. *α*2 adrenergic agonist at high concentrations blocks the release of norepinephrine from neurons projecting from LC to the extended areas, including the thalamus, the cortex and the arousal pathways, resulting in wide-spread slow-delta oscillations, consistent with the dynamics shown in the unresponsive state in our study. When the *α*2 adrenergic agonist concentration is slowly lowering following the end of anesthetic infusion, the blockade of norepinephrine release may be limited to the preoptic area, which activates inhibitory GABAergic projections in the arousal centers (Zhang *et al.*, 2015), similar effects of GABAergic general anesthetic agents. Increasing GABA_A_ conductance leads to synchronous alpha-activity in the thalamocortical loops (Ching *et al.*, 2010), consistent with our finding of inter-regionally coherent alpha oscillations during the intermediate state in early recovery from dexmedetomidine.

In addition to the elevated spindle activity during early recovery, we found that a transient increase in the spindle activity at behavioral transitions. The spindles are thought to increase an arousability threshold since burst firing of thalamocortical cells during spindles blocks external stimuli and possibly protects sleep against the stimuli (Wimmer *et al.*, 2012; Astori *et al.*, 2013). Spindles during sleep have also been reported to increase at transitional periods out of NREM sleep (Prerau *et al.*, 2017). Characteristic features of the spindle activities with dexmedetomidine seem to closely approximate the spindles during NREM sleep, suggesting common mechanisms (Takeuchi *et al.*, 2016). Dexmedetomidine is shown to activate endogenous NREM sleep-promoting pathways (Nelson *et al.*, 2003; Yu *et al.*, 2018). Clinically, these spindles can be useful as a sign for state changes under dexmedetomidine. When the patients are anesthetized under dexmedetomidine with minimum spindle activity, monitoring the spindle activity can be preventive against lightening the anesthetic level. When conscious or arousable sedation is appropriate, constant appearance of the high spindle activity may be a goal. Human EEG under dexmedetomidine shows dominant slow-delta oscillations and alpha-theta oscillations following LOC and the spindles in the frontal region (Akeju *et al.*, 2014b; Akeju *et al.*, 2016), consistent with our LFP findings. Together, forehead application of the EEG monitors can be modified for this new role of monitoring spindles under dexmedetomidine sedation and anesthesia.

We demonstrate for the first time that the use of an *α*2 antagonist instantly restores the highest level of task performance and concurrent awake network dynamics. The 3D state space dynamics clearly characterizes a discontinuous transition upon antagonist administration without transitioning through intermediate states. During emergence without antagonist, we found diffuse clouds or possible clusters connecting between unresponsiveness and a performing state shown in **Figure 6**. This intermediate state corresponds to spectrographic signatures, such as an increase in the alpha power and the spindle activities. Hudson and colleagues first reported multiple metastable states during recovery from isoflurane anesthesia in rodents (Hudson *et al.*, 2014). Proekt and Hudson further delineated stochastic dynamics of neuronal states during anesthetic recovery and explained that anesthetic recovery can be independent from pharmacokinetics (Proekt and Hudson, 2018). Our results with dexmedetomidine are consistent with the proposed multi-state stochastic recovery theories. However, our results demonstrate that the probabilistic process of anesthetic recovery can be shut down by an overwhelming pharmacological effect of *α*2 adrenergic antagonist at the receptor level and that the intermediate recovery states can be totally bypassed. Moreover, our results support that the brain is capable to switch dynamics in and on-and-off manner and the expected behavioral performance can accompany without delay.

How the unconscious state can be actively reversed has been investigated in anesthetized subjects and subjects with coma due to other causes with a limited degree of recovery (Alkire *et al.*, 2007; Schiff *et al.*, 2007; Xu *et al.*, 2019). Gao and colleagues recently reported that activating nucleus gigantocellularis neurons in the reticular activating system elicited a high degree of awakening, including cortical, autonomic, and behavioral recovery, in both isoflurane anesthetized and hypoglycemia-induced coma rodents (Gao *et al.*, 2019). Activation of dopamine and norepinephrine pathways are also shown to promote behavioral arousal in anesthetized rodents with inhaled anesthetics (Kenny *et al.*, 2015; Taylor *et al.*, 2016). In these animal studies, the level of performance recovery is not yet clear. Clinically, emergence and postoperative neurocognitive problems still affect patients undergoing surgery of all ages (Lepouse *et al.*, 2006; Vlajkovic and Sindjelic, 2007; Monk and Price, 2011) and are thought to be associated with an overall increase in morbidity and mortality (Monk *et al.*, 2008; Steinmetz *et al.*, 2009; Witlox *et al.*, 2010). Immediate restoration of the top task performance by an *α*2 adrenergic antagonist in the current study is promising and suggests its future clinical role in active awakening. Neurocognitive problems that may occur during anesthetic emergence can be avoidable by appropriate antagonism of the anesthetic action.

In summary, behavioral endpoints, such as LOC, ROC and full task performance recovery, ROPAP, are successfully defined during *α*2 adrenergic agonist-induced altered states of consciousness in non-human primates, and are all associated with an distinctive change in the cortical dynamics, consistent with the abrupt state transitions during propofol anesthesia and recovery (Ishizawa *et al.*, 2016; Patel *et al.*, 2020). The intermediate recovery state between unresponsiveness and full performance recovery is distinguished by sustained high spindle activities with rapidly fluctuating behavioral responses. Awakening by *α*2 adrenergic antagonist completely eliminates the intermediate state and discontinuously restores awake dynamics and the top task performance while dexmedetomidine was still being infused. The results suggest that instant functional recovery is possible following anesthetic-induced unconsciousness and the intermediate state is not a necessary path for the brain recovery.

## Materials and methods

All animals were handled according to the institutional standards of the National Institute of Health. Animal protocol was approved by the institutional animal care and use committee at the Massachusetts General Hospital (2006N000174). We used two adult male monkeys (*macaca mulatta*, 10 - 12 kg). Prior to starting the study, a titanium head post was surgically implanted on each animal. A vascular access port was also surgically implanted in the internal jugular vein (Model CP6, Access Technologies). Once the animals had mastered the following task, prior to the recording studies, extracellular microelectrode arrays (Floating Microelectrode Arrays, MicroProbes) were implanted into the primary somatosensory cortex (S1), the secondary somatosensory cortex (S2) and ventral premotor area (PMv) through a craniotomy (**Fig. 1A**). Each array (1.95 × 2.5 mm) contained 16 platinum-iridium recording microelectrodes (~0.5 Meg Ohm, 1.5 - 4.5 mm staggered length) separated by 400 μm. The placement of arrays was guided by the landmarks on cortical surface (**Fig. 1A**) and stereotaxic coordinates (Saleem and Logothetis, 2012). A total of five arrays were implanted in Monkey 1 (2 arrays in S1, 1 in S2, and 2 in PMv in the left hemisphere) and four arrays in Monkey 2 (2 arrays in S1, 1 in S2, and 1 and PMv in the right hemisphere). Data recorded from S2 arrays were not included in this study. The recording experiments were performed after 2 weeks of recovery following the array surgery. All experiments were conducted in the radio-frequency shielded recording encolsures.

The animals were trained in the behavioral task shown in **Figure 1B**. After the start tone (1000 Hz, 100 msec) the animals were required to initiate each trial by holding the button located in front of the primate chair using the hand ipsilateral to the recording hemisphere. They were required to keep holding the button until the task end in order to receive a liquid reward. The monkeys were trained to perform correct response greater than 90% of the trials consistently for longer than ~1.5 hours in an alert condition. Animal’s performance during the session was monitored and simultaneously recorded using a MATLAB based behavior control system (Asaad and Eskandar, 2008a, b). The animal’s trial-by-trial button-holding responses were used to define two metrics to allow quantification of behavioral endpoints: task engagement and task performance probability (Patel *et al.*, 2020). Task engagement indicates a probability of any response initiation, including correct responses and failed attempts, and task performance represents a probability of correct responses only (Wong *et al.*, 2011; Wong *et al.*, 2014) (**Fig. 1B, C**). LOC was defined as the time at which the probability of task engagement was decreased to less than 0.3, and ROC was defined as the first time, since being unconscious, at which the probability of task engagement was greater than 0.3 (Chemali *et al.*, 2011; Mukamel *et al.*, 2014) (**Fig. 1C**). We also defined return of preanesthetic performance level (ROPAP) at which the probability of task performance was returning to greater than 0.9 since being unconscious and remained so for at least 3 minutes.

Dexmedetomidine was infused for total 60 minutes at 18 μg/kg/h for the first 10 minutes and then 4 μg/kg/h for 50 minutes through a vascular access port. The infusion rate of dexmedetomidine was determined in order to induce LOC in approximately 10 minutes in each animal. α2 adrenergic antagonist atipamezole (100 μg/kg) was injected through a vascular access port at 30 minutes of dexmedetomidine infusion while it was still being infused in 4 sessions and at the end of 60-min dexmedetomidine infusion in one session. No other sedatives or anesthetics were used during the experiment. The animal’s heart rate and oxygen saturation were continuously monitored throughout the session (CANL-425SV-A Pulse Oximeter, Med Associates). The animals maintained greater than 94% of oxygen saturation throughout the experiments.

Neural activity was recorded continuously and simultaneously from S1, S2, and PMv through the microelectrode arrays while the animals were alert and participating in the task and throughout anesthesia and recovery. Analog data were amplified, band-pass filtered between 0.5 Hz and 8 kHz and sampled at 40 kHz (OmniPlex, Plexon). Local field potentials (LFPs) were separated by low-pass filtering at 200 Hz and down-sampled at 1 kHz. The spiking activity was obtained by high-pass filtering at 300 Hz, and a minimum threshold of three standard deviations was applied to exclude background noise from the raw voltage tracings on each channel. Action potentials were sorted using waveform principal component analysis (Offline Sorter, Plexon). The recordings under dexmedetomidine anesthesia were performed 8 times in Monkey 1 and 9 times in Monkey 2. In separate sessions, the recordings were performed under dexmedetomidine in the animals that were required no task performance (2 sessions in Monkey 1 and 2 sessions in Monkey 2) and in the blind-folded animals that were performing the task (2 sessions in Monkey 2). Arousability was tested in 2 sessions in Monkey 2. In addition, multiple recordings were performed in alert performing animals without aneshtesia.

All LFP analyses were performed using existing and custom-written functions in MATLAB (MathWorks Inc., Natick, MA). Raw signals were filtered to remove 60 Hz noise by running a 2^nd^ order Butterworth filter in each time direction. The resulting signals were subject to continuous multitaper spectral and channel-to-channel coherence analysis using the Chronux toolbox (Mitra and Bokil, 2008). To generate the spectrograms, full-length session were processed by running a thirty second, non-overlapping, time window, applying three tapers. The spectral lines show the average power of all channels, and 95% confidence intervals, from one-minute epoch of each condition analyzed by Welch’s power spectral density estimate. Statistical analyses on the spectral change were performed by comparing the awake values, frequency by frequency, versus other conditions using ANOVA test and *post-hoc* Bonferroni multiple comparisons with the significance threshold set to 0.05. Channel-to-channel coherence calculations were performed running a two second, non-overlapping window. To generate coherograms we averaged all pairwise combinations within or between arrays. The average coherence for each condition was computed from 30 consecutive windows (one minute) within each condition. Statistical comparisons of local and inter-regional coherence between the different conditions were performed using ANOVA test and *post hoc* Bonferroni multiple comparisons test with the significance threshold set to 0.05.

Spindle detection and characterization was addressed using custom-written functions in MATLAB. We based our analysis on the methodology described by Kam and colleagues (Kam *et al.*, 2019) using the FMA toolbox (Khodagholy *et al.*, 2017). Briefly, we bandpass-filtered the 60 Hz noise-free LFP traces in the 9-17 Hz range. Then, we located spindle-like events by detecting their start/end (filtered signal amplitude threshold set at 4 z-scores above the baseline) and peak (threshold set at 6 z-scores). Only events with a total duration between 500 and 2500 milliseconds were further analyzed. After visual confirmation and removal of artefacts (events with peak amplitudes >15 z-scores), we counted events located within each periods of interest: the last ten minutes before the anesthesia infusion started (awake), ten minutes under the effects of anesthesia (anesthesia) and ten minutes during performing period after ROC (recovery) or after the infusion of alpha-antagonist started (antagonist). We calculated the spindle density as number of events per minute during each period of time. The spindle peak frequency was calculated for each event’s raw signal spectrogram using the Hilbert-Huang transform (Huang *et al.*, 2016) and obtaining their frequency of maximum power.

To characterize the dexmedetomidine-induced brain states and their transitions, a three-dimensional space was defined using three spectral power ratios (Gervasoni *et al.*, 2004; Patel *et al.*, 2020). Briefly, we obtained three frequency-band spectral power ratios (Ratio1 = power_17-35 Hz_/power_0.5-60 Hz_, Ratio2 = power_1-8 Hz_/power_1-16 Hz_ and Ratio3 = power_9-15Hz_/power_1-16 Hz_) from each channel. For each region and ratio, data were concatenated in a time-by-channel matrix and subjected to principal component analysis (PCA). The first principal component (PC1) explained more than 70% of the data’s variance. To reduce instant variability, we smoothed the PC1 values by running a 20-sec Hanning window. All figures are scatter representations of the three ratios’ PC1s where each point represents one second of time. The data density was calculated with a kernel density estimator function. The speed was calculated as the Euclidean length between consecutive points. The trajectory-speed plots show the speed values only for 20 minutes of data around the LOC, ROC or antagonist administration. The engagement plots represent the same values of probability calculated above.

## Acknowledgements

We thank Emad N. Eskandar, MD for performing craniotomy for electrodes implantation, Tatsuo Kawai, MD for performing vascular port surgeries, Shaun R. Patel, PhD for providing training for the data analyses, and Warren M. Zapol, MD for guiding and supporting the project development.

## Author Contributions

Y.I. designed the study and performed the experiments. J.J.B., J.B. and Y.I. analyzed the data. Y.I. wrote the manuscript. All authors edited the manuscript.

